# Tetrahydrobiopterin restores oligodendrocyte maturation after hypoxic white matter injury by reprogramming translational output

**DOI:** 10.64898/2026.10.23.761441

**Authors:** Nemanja Sarić, Ingrid Antonios, Karl Heiser, Alissa Ballard, Kennedy Miller, Iman Mansoor, May T Rajtboriraks, Shreya Chandran, Evan Goldstein, Nobuyuki Ishibashi

## Abstract

**Background:** Complex congenital heart defects (CHDs) are frequently associated with chronic cerebral hypoxemia, leading to impaired white matter maturation. One proposed mechanism involves hypoxia-induced disruption of the nitric oxide synthase cofactor tetrahydrobiopterin (BH4), resulting in oxidative stress and impaired oligodendrocyte (OL) development. While our previous studies found BH4 supplementation improves hypoxia-induced white matter dysmaturation, the molecular mechanisms remain incompletely understood. Because OL maturation requires coordinated translational control, we hypothesized that BH4 promotes white matter maturation by restoring translation programs disrupted by hypoxia.

**Methods:** Translating Ribosome Affinity Purification sequencing (TRAP-seq) was performed on OL lineage cells isolated from transgenic ribosomal reporter mice exposed to chronic hypoxia (10.5% O₂, postnatal day 3–11) with or without daily BH4 treatment. Differential translation, weighted gene co-expression network analysis, pathway enrichment, RNAscope, and immunohistochemistry identified translational programs associated with hypoxia and BH4-mediated rescue.

**Results:** Developmental hypoxia extensively altered the OL translatome, suppressing transcripts associated with OL maturation while enriching pathways related to translational regulation and cellular stress. Network analyses identified BH4-responsive gene modules enriched for OL maturation, protein metabolism, and translation-related processes. BH4 partially reversed hypoxia-induced translational changes, restored OL maturation gene expression, and normalized nuclear localization of the lineage regulator OLIG1, consistent with recovery of maturation-associated regulatory programs.

**Conclusions:** Developmental hypoxia disrupts OL maturation through coordinated alterations in ribosome-associated gene expression and post-transcriptional regulatory pathways. BH4 promotes recovery of OL maturation programs, including restoration of translation-related networks and OLIG1 nuclear localization, providing mechanistic insight into BH4-supported white matter maturation under chronic hypoxic conditions.

**Research Perspective:** *What New Question Does This Study Raise?:* Developmental hypoxemia in congenital heart disease remodels the oligodendrocyte translatome, raising the question of whether translational dysregulation represents a fundamental and therapeutically targetable mechanism underlying white matter dysmaturation in other forms of perinatal brain injury.

*What Question Should Be Addressed Next?:* Future studies should determine how BH4 restores ribosome-associated translation of oligodendrocyte maturation programs, establish whether these molecular changes increase myelin protein synthesis and improve long-term neurodevelopmental outcomes, and define the cross-species efficacy for clinical translation.

*Clinical Research Implication:* These findings provide a mechanistic rationale for evaluating BH4 as a neuroprotective adjunct therapy to preserve white matter development in fetuses and infants with congenital heart disease experiencing chronic cerebral hypoxemia.

## INTRODUCTION

Congenital heart defects (CHDs) represent the most prevalent live birth defects diagnosed in newborns in the United States, with recent estimates pointing to an incidence of ∼1% of births^1–3^. Children with complex CHDs frequently experience chronic cerebral hypoxemia beginning in utero and persisting through the perinatal period due to impaired cardiac output and altered cerebrovascular regulation^4–6^. These sustained reductions in cerebral oxygen delivery are closely linked to abnormalities in white matter development, including diffuse hypomyelination and arrested oligodendrocyte (OL) maturation, which are prominent contributors to the neurodevelopmental deficits observed in children with CHD^7–11^.

A growing body of work implicates oxidative stress as a central mechanism by which hypoxia disrupts and stalls normal OL lineage progression^12,13^. One key pathway involves the oxidation-driven depletion of the nitric oxide synthase (NOS) cofactor tetrahydrobiopterin (BH4), which leads to NOS uncoupling, impaired nitric oxide signaling, and exacerbation of reactive oxygen species (ROS) production^14–16^. This oxidative microenvironment disrupts signaling pathways that govern OL maturation and myelin production while increasing the metabolic burden placed on developing OLs, whose maturation requires coordinated protein synthesis to support rapid membrane expansion and myelin production. OL maturation and myelination are among the most biosynthetically demanding processes in the developing central nervous system, requiring coordinated synthesis of proteins involved in lineage progression, membrane expansion, lipid metabolism, and myelin formation. Consequently, immature OLs are particularly vulnerable to disturbances in cellular redox homeostasis and energy metabolism^17^. Consistent with this concept, our previous work demonstrated that BH4 supplementation during developmental hypoxia restores OL maturation, improves myelination, and ameliorates neurobehavioral deficits, identifying BH4 bioavailability as a critical determinant of white matter resilience under hypoxic conditions^18^. However, the mechanisms through which BH4 confers protection to OL lineage cells and promotes their maturation remain poorly understood.

Emerging evidence indicates that oxidative and metabolic stress regulate cell fate not only through transcriptional changes but also by remodeling mRNA translation. Stress-responsive pathways, including the integrated stress response (ISR) and mTOR signaling, suppress canonical translation initiation while selectively maintaining ribosome association of transcripts that promote stress adaptation and cellular survival^19–21^. As a result, translational output can be redistributed among functionally distinct mRNA populations rather than uniformly reduced. This process, often termed translational selectivity or translational reprogramming, allows stressed cells to prioritize synthesis of proteins required for immediate survival at the expense of lineage-specific maturation programs. Such selective translational regulation may be especially important during OL maturation, when extensive proteome remodeling is required to support membrane expansion, myelin protein synthesis, and myelin sheath formation. Whether developmental hypoxia alters ribosome-associated gene expression programs in OL lineage cells, and whether restoration of BH4 rebalances these translational programs toward maturation, has not been investigated.

To define the translational mechanisms through which BH4 supports white matter development, we combined a well-established mouse model of developmental hypoxia with Translating Ribosome Affinity Purification sequencing (TRAP-seq) to profile ribosome-associated mRNAs within postmitotic oligodendrocytes. We demonstrate that developmental hypoxia extensively remodels the postmitotic OL translatome by reciprocally regulating ribosome-associated gene networks with overlapping biological functions, consistent with selective translational reprogramming rather than global translational suppression. BH4 rebalances these translational programs by restoring ribosome association of key OL maturation transcripts while reducing ribosome association of hypoxia-induced translation initiation transcripts. This translational reprogramming is accompanied by recovery of OLIG1 protein expression and nuclear localization, supporting a mechanism whereby BH4 promotes OL maturation through restoration of lineage-specific translational programs under chronic hypoxic conditions.

## METHODS

### Animals

C57BL/6J (BL6) and *CNP-bacTRAP* (JAX strain #009159) mice were bred and kept at the Children’s National Comparative Medicine Unit (CMU) and Boston University Animal Science Center (ASC) using standard housing conditions. Pups from both sexes were included in the hypoxia exposure experiments at P3, and litters were randomly allocated to one of the studied groups. The study was conducted in compliance with the NIH Guide for the Care and Use of Laboratory Animals. Approval was obtained by the Animal Care and Use Committees of Children’s National Hospital and Boston University.

### Hypoxia exposure and treatments

The hypoxic exposure period extended from postnatal day 3-11 (P3-P11), equivalent to the third trimester of human pregnancy^13^. Pups from *CNP-bacTRAP* or BL6 crosses were fostered at P1-P2 to CD-1 strain dams for maintaining optimal nutrition and maternal care. Cages with dams and litters were placed in a hypoxic chamber system (Biospherix, Redfield, NY) with a constant oxygen level of 10.5% ± 0.5 (Pro Ox model 360 Biospherix). Nitrogen gas was used to displace oxygen and maintain hypoxic conditions within the chamber. Temperature and humidity were kept comparable to normoxic conditions. Age-matched pups were used as controls and remained with their mother in normoxic conditions for the equivalent study period.

As performed in our previous studies^18^, CD-1 dams in all BH4 treatment groups received synthetic BH4 (Sapropterin Dihydrochloride powder for oral solution, PAR pharmaceutical, 49884-0948-72) through oral gavage at a dose of 50 mg/kg/day from P3-P11, thus modeling initiation of therapy at the beginning of the third trimester -a common time for CHD diagnosis. BH4 in powder form was diluted in normal saline at a 10 mg/ml concentration. A total volume of 150 µl/day was administered through a flexible feeding tube (Instech Laboratories Inc, FTP-20-38). Control groups received the same volume of saline alone (VEH) through oral gavage over the same period.

### Tissue collection for histology

On P11, anesthesia was induced with a mix of ketamine (100 mg/kg) and xylazine (20 ug/kg), injected intraperitoneally. Pups were perfused with ice cold intracardiac PBS followed by tissue fixation with 4% paraformaldehyde (PFA). The brains were then kept in 4% PFA overnight at 4 °C, followed by a sucrose gradient (15-30%) successively for 24 hours each. The tissues were subsequently flash frozen in optimal cutting temperature (OCT) medium and stored at –80 °C. For preparing frozen sections, brains were coronally sliced using a cryostat (Thermo Scientific Cryostat Microm HM 525) into 20 µm sections and mounted on microscope slides (Superfrost plus microscope slides, Fisher brand, 22037246). Slides were stored at –80 °C prior to staining.

### TRAP RNA sequencing

RNA was isolated from *CNP-bacTRAP* mice (*n =* 4−5 per group) using a modified version of the previously described protocol^22,23^. Subcortical white matter was dissected in ice-cold dissection buffer (2.5 mM 4-(2-hydroxyethyl)-1-piperazineethanesulfonic acid (HEPES), 35 mM glucose and 4 mM NaHCO_3_, and 100 μg/ml cycloheximide in Hank’s balanced salt solution (HBSS)), homogenized in lysis buffer (20 mM HEPES, 150 mM KCl and 10 mM MgCl_2_, 0.5 mM dithiothreitol (DTT), 100 μg/ml cycloheximide, 10 μl/ml rRNasin, 10 μl/ml Superasin, and ethylenediaminetetraacetic acid (EDTA)-free protease inhibitors in RNase-free water), and then centrifuged at 1000 × *g* for 10 min. 1% NP-40 and 1% DHPC were added to the supernatants, chilled on ice for 5 min, and centrifuged at 10,000 × *g* for 10 min. Supernatants were incubated overnight at 4 °C with GFP antibodies (HtzGFP-19F7 and HtzGFP-19C8, Memorial Sloan Kettering Center) bound to Streptavidin MyOne T1 Dynabeads (Invitrogen). The beads were then washed four times in a high salt buffer (20 mM HEPES, 350 mM KCl, 10 mM MgCl_2_, 1% NP-40, 0.5 mM DTT, and 100 μg/ml cycloheximide in RNase-free water), and RNA was isolated using the Absolutely RNA Nanoprep Kit (Agilent). RNA samples were sent to the Genome Sciences and Bioinformatics core at Penn State College of Medicine for Library preparation and RNA-sequencing. RNA quality was determined using the Bioanalyzer Pico Kit (Agilent), and quantity was determined with the Qubit fluorometer (ThermoFisher). Libraries were prepared from RNA (RIN > 7) using the Trio RNA-Seq Library Preparation Kit (NuGen), according to the manufacturer’s protocol. Libraries were sequenced using the NovaSeq 6000 System (Illumina) to obtain 50 base pair (bp), paired-end reads. The first 5 bp were trimmed from the 5′ end of the reads using Trimmomatic^24^, as recommended by the Library Preparation Kit. Reads were mapped to the mouse reference genome (Genome Reference Consortium Mouse Build 38 – mm10) using STAR^25^, and read counts were generated using HTseq^26^.

Raw read counts were imported into DESeq2 (v1.40.2) in R for normalization, dispersion estimation, and differential enrichment testing using the Wald method with Benjamini–Hochberg correction for multiple comparisons. Genes were considered significantly differentially translated at an adjusted p-value (padj) < 0.05 and an absolute log₂ fold change (|log₂FC|) ≥ 0.25, unless otherwise specified. For the analysis of transcript translational rescue, genes were defined as “rescued” if their transcript translation was significantly downregulated in HX+VEH vs NX+VEH (padj < 0.05, |log₂FC| ≥ 0.25) and significantly upregulated in HX+BH4 vs HX+VEH comparisons under the same criteria. Normalized counts and differential enrichment statistics were used for downstream analyses of translational profiles. Gene Ontology (GO) and pathway enrichment analyses were performed using Enrichr (https://maayanlab.cloud/Enrichr/) and/or KEGG. Lists of significantly up-or downregulated genes were submitted separately, and enrichments for biological process, molecular function, cellular component, and KEGG pathway terms were assessed. Enrichr adjusted p values were used for ranking and visualization. Selected GO categories were visualized using bar plots and network representations generated in R or Python. Weighted gene co-expression network analysis was performed using the WGCNA package in Python to identify coordinated patterns of ribosome-associated gene expression across experimental conditions. Normalized expression values from DESeq2 were used as input for network construction. Genes with low enrichment across samples were filtered prior to analysis. An adjacency matrix was generated using a soft-thresholding power selected to approximate scale-free network topology, followed by calculation of the topological overlap matrix and hierarchical clustering. Modules of co-expressed genes were identified using dynamic tree cutting and module merging based on eigengene similarity. Module eigengenes were correlated with experimental conditions to identify modules associated with developmental hypoxia and BH4 treatment. Genes within significant modules were further analyzed using transcription factor enrichment and pathway enrichment approaches.

### RNAscope staining

RNAscope staining was performed using standard multiplex fluorescent staining protocols provided by Advanced Cell Diagnostics (ACD). Briefly, slides were taken out of the freezer and kept at room temperature for 15 minutes, followed by warming at 37 °C for 20 minutes. OCT was removed by immersing slides in PBS for 5 minutes with agitation. Slides were then left to bake at 60 °C for 30 minutes to ensure adhesion. Once ready to use, slides were dehydrated by immersion in successive ethanol solutions at increasing concentrations (50%,70%,100%,100%) for 5 minutes each. Hydrophobic barriers were then drawn around the sections using an ImmEdge Pen (Vector Laboratories, H-4000). Sections were first covered with hydrogen peroxide (H_2_O_2_) and incubated for 10 minutes at room temperature followed by protease III with an incubation period of 30 minutes in a HybEZ II Oven (HybEZ™ II Hybridization System for Manual Assays, PN 321710/321720) at 40 °C. Following this, sections were incubated with RNA probes (*Olig2*, *Olig1*) that were pre-warmed for 10 min at 40 °C. Channel 1 probes were used at the manufactured concentration. Channel 2 probes were diluted in TSA buffer to a target concentration of 1:500. Slides were incubated for 2 hours at 40 °C then stored overnight in 5x saline-sodium citrate (SSC) at room temperature.

On the second day, signal amplification with Amp 1, 2 and 3 mixes was performed using recommended incubation periods (30 minutes, 30 minutes and 15 minutes respectively) in the HybEZII Oven at 40 °C. Sections were then covered with HRP-C1 (or C2 based on channel being used) and incubated at 40 °C for 15 minutes. Fluorophore solutions containing Opal 570 or 690 (depending on probe channel) (Opal 570/690 reagent pack, Akoya Biosciences) diluted in TSA buffer at a target concentration of 1:500 were then incubated for 30 minutes at 40 °C. Finally, signal development was terminated using HRP blocker incubation for 15 minutes at 40 °C. Coverslips were mounted using DAPI-containing mounting medium (DAPI fluoromount–G, SouthernBiotech, 0100-20), and the slides were left at room temperature, in the dark, overnight for curing prior to imaging.

### Confocal imaging

Confocal images of RNAscope staining (*Olig1*+*Olig2*) were obtained using Nikon A-1 HD25 and Evident FV4000 confocal microscopes. High resolution (1024 x 1024 pixels) z series scans with a 0.5 μm z step interval and consisting of 35-50 optical planes were taken for *Olig1*+*Olig2* stained slides with a 20x air objective and 3x digital zoom. Three regions of interest (ROIs) from 3-4 slices/animal were acquired from white matter regions encompassing the corpus callosum, external capsule and cingulum for downstream image analysis.

### Image analysis

Confocal image analyses were performed using Imaris (v10.2 and v11; Oxford Instruments) and ImageJ/Fiji. For RNAscope analyses, OL lineage cells were first segmented based on OLIG2 immunoreactivity by a blinded investigator using the Imaris surface segmentation function. RNAscope puncta were subsequently segmented using the surface function, and colocalization with OLIG2-positive cells was determined prior to quantification. *Olig1* mRNA expression was quantified as the mean RNAscope signal intensity within segmented OLIG2-positive cells. For assessment of OLIG1 protein localization, OLIG1-positive OL lineage cells were manually quantified in ImageJ/Fiji by a blinded investigator and classified as exhibiting either predominantly nuclear or predominantly cytoplasmic localization based on the distribution of OLIG1 immunofluorescence relative to the DAPI-defined nucleus.

### Statistical analysis

All statistical analyses were performed using GraphPad Prism v10 (GraphPad Software, San Diego, CA) unless otherwise indicated. Data are presented as mean ± SEM, with individual animals serving as the experimental unit (n) for all analyses. Image acquisition and quantitative image analyses were performed blinded to experimental group assignment. Data distributions were assessed for normality using the Anderson–Darling, D’Agostino–Pearson omnibus, and Shapiro–Wilk tests. For normally distributed datasets, comparisons among experimental groups were performed using one-way analysis of variance (ANOVA) followed by Tukey’s multiple-comparisons test.

For datasets that did not meet assumptions of normality, comparisons were performed using the Kruskal–Wallis test followed by Dunn’s multiple-comparisons test. All statistical tests were two-sided, and P < 0.05 was considered statistically significant. Differential ribosome-associated gene expression from TRAP-seq was analyzed using DESeq2 with Wald tests, applying Benjamini–Hochberg false discovery rate correction for multiple testing. Differentially translated genes were defined as those with an absolute log2 fold change ≥ 0.25 and adjusted P < 0.05. Weighted gene co-expression network analysis (WGCNA) module eigengenes were correlated with experimental groups using Pearson correlation coefficients. Functional enrichment analyses, including Gene Ontology, KEGG, WikiPathways, ClinVar, ChEA, and STRING analyses, were performed using the respective analytical platforms, with significance determined using their reported multiple testing-adjusted P values.

## RESULTS

### BH4 preferentially restores hypoxia-suppressed ribosome-associated transcripts in OL lineage cells

To define how developmental hypoxia alters ribosome-associated gene expression within the OL lineage and determine whether BH4 supplementation restores these changes, we performed Translating Ribosome Affinity Purification sequencing (TRAP-seq) using *CNP-bacTRAP* mice (Figure 1A, B). Ribosome-associated mRNAs were isolated from postmitotic OLs at postnatal day 11 following chronic normoxia (NX/Veh), hypoxia (HX/Veh), or hypoxia with daily BH4 treatment (HX/BH4), enabling cell type-specific profiling of the actively translated transcriptome (Figure 1C).

**Figure 1.**
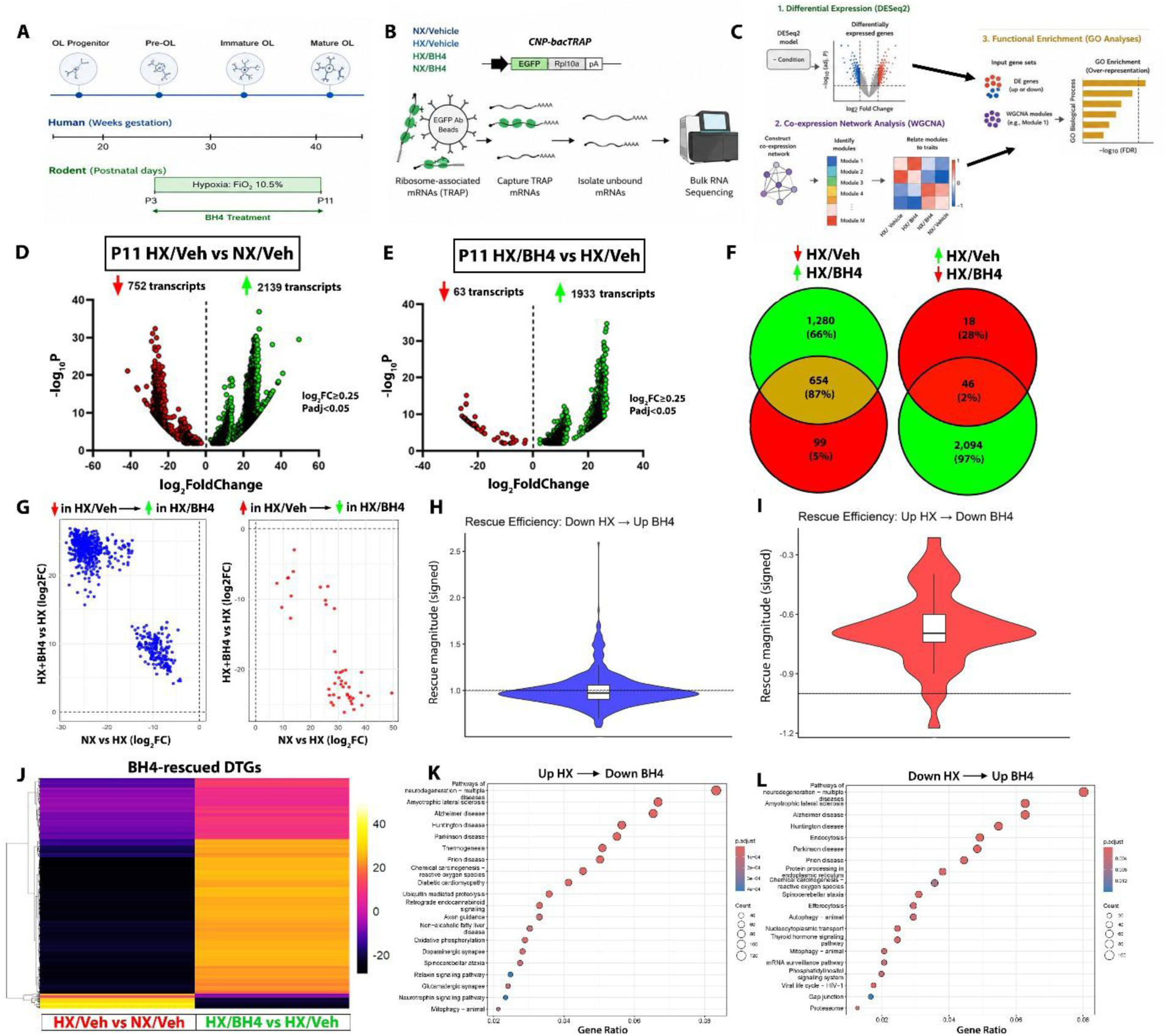
Hypoxia remodels the OL translatome and BH4 restores a substantial subset of dysregulated translational programs. (A) Developmental timeline illustrating OL (OL) lineage progression from progenitors to mature OLs, the developmental window corresponding to rodent postnatal days P3–P11, and the chronic hypoxia (HX; 10.5% O₂) exposure paradigm with daily BH4 administration. Human developmental equivalence and timing of white matter vulnerability are indicated. (B) Experimental workflow. *CNP-bacTRAP* mice were used to isolate ribosome-associated mRNAs from OL lineage cells under normoxia with vehicle (NX/Veh), hypoxia with vehicle (HX/Veh), and hypoxia with BH4 treatment (HX/BH4). Purified ribosome-bound RNAs underwent bulk RNA sequencing followed by differential translation, weighted gene co-expression network analysis (WGCNA), and transcript feature analyses. (C) Bioinformatic analysis pipeline summarizing differential translation (DESeq2), co-expression network analysis (WGCNA), transcript feature characterization, and downstream pathway enrichment analyses used to identify BH4-responsive translational programs. (D) Volcano plot showing differentially translated genes (DTGs) in HX/Veh versus NX/Veh at P11. Red points indicate transcripts significantly reduced by hypoxia and green points indicate transcripts significantly increased by hypoxia (|log₂FC| ≥ 0.25, adjusted P < 0.05). Numbers denote significantly altered transcripts. (E) Volcano plot of DTGs in HX/BH4 versus HX/Veh. Green points represent transcripts increased following BH4 treatment, whereas red points represent transcripts reduced by BH4 relative to hypoxic controls. (F) Venn diagrams illustrating overlap between hypoxia-responsive DTGs and BH4-responsive DTGs. Left, transcripts downregulated by hypoxia and restored by BH4. Right, transcripts upregulated by hypoxia and reduced by BH4. Central intersections represent BH4-rescued translational targets. (G) Scatterplots comparing hypoxia-induced translational changes with BH4-mediated responses for transcripts decreased (left) or increased (right) during hypoxia. Each point represents an individual DTG. (H–I) Violin plots summarizing rescue efficiency for transcripts downregulated by hypoxia and increased by BH4 (H) or upregulated by hypoxia and reduced by BH4 (I). Rescue efficiency was calculated as the BH4-induced change relative to the magnitude of the hypoxia-induced change. Embedded boxplots indicate median and interquartile range. (J) Heatmap of BH4-rescued DTGs demonstrating opposing translational responses across HX/Veh versus NX/Veh and HX/BH4 versus HX/Veh comparisons. Rows represent individual genes and columns represent experimental comparisons. (K–L) KEGG enrichment analyses of BH4-rescued transcripts. Panel K shows biological processes enriched among transcripts increased by BH4 following hypoxia, whereas panel L summarizes pathways enriched among transcripts decreased by BH4. Dot size represents gene count and color indicates statistical significance. Differential translation statistics were derived from DESeq2 analyses using Wald tests with Benjamini–Hochberg correction for multiple comparisons (adjusted p < 0.05).

Developmental hypoxia produced widespread remodeling of the OL translatome (Figure 1D). Using a threshold of |log₂FC| ≥ 0.25 with Benjamini–Hochberg adjusted P < 0.05, we identified 2,891 differentially translated genes (DTGs) between HX/Veh and NX/Veh, including 752 transcripts exhibiting reduced ribosome association and 2,139 transcripts exhibiting increased ribosome association during hypoxia. Comparison of HX/BH4 and HX/Veh identified 1,996 BH4-responsive DTGs, the majority of which (1,933 transcripts) demonstrated increased ribosome association following BH4 treatment, whereas only 63 transcripts were reduced (Figure 1E). These findings suggest that BH4 preferentially restores ribosome association among transcripts suppressed during developmental hypoxia.

To determine the extent to which BH4 reversed hypoxia-induced translational changes, we compared DTGs identified in the hypoxia and BH4 datasets. Notably, among transcripts reduced during hypoxia, 87% (654) of the BH4-responsive genes overlapped with the hypoxia-downregulated dataset, representing a large majority of all hypoxia-suppressed transcripts (Figure 1F, left). In contrast, only 46 transcripts overlapped between hypoxia-upregulated and BH4-downregulated gene sets (Figure 1F, right), indicating that BH4 exerts a substantially greater restorative effect on transcripts whose ribosome association is diminished by hypoxia than on transcripts elevated during hypoxic stress.

Scatterplot analyses further illustrated the directional relationship between hypoxia-and BH4-induced translational changes (Figure 1G). Transcripts suppressed during hypoxia generally exhibited increased ribosome association following BH4 treatment, whereas transcripts elevated during hypoxia showed comparatively modest reductions with BH4. Quantification of rescue efficiency confirmed this asymmetry (Figure 1H, I). Transcripts reduced by hypoxia frequently approached or exceeded complete restoration following BH4 administration, whereas the corresponding rescue efficiency for hypoxia-induced transcripts was substantially lower, consistent with preferential recovery of developmentally important translational programs rather than broad suppression of hypoxia-responsive translation.

Hierarchical clustering of BH4-rescued DTGs demonstrated coordinated restoration of ribosome-associated transcript abundance across experimental groups (Figure 1J). To further define the biological processes represented within these rescued transcript sets, KEGG pathway enrichment analyses were performed separately for transcripts reduced by hypoxia and restored by BH4, and for transcripts increased by hypoxia and reduced following BH4 treatment. Surprisingly, both gene sets exhibited strong enrichment for overlapping neurodegeneration-associated pathways, including Alzheimer’s disease, Huntington’s disease, Parkinson’s disease, amyotrophic lateral sclerosis, and prion disease (Figure 1K, L). Although these pathway annotations were shared, they arose from distinct subsets of reciprocally regulated transcripts, suggesting that developmental hypoxia and BH4 treatment remodel translation within common biological systems rather than uniformly altering entire pathways.

To determine whether BH4-mediated translational remodeling persisted beyond the period of active OL maturation, we performed TRAP-seq analysis at postnatal day 30 (P30). In contrast to the extensive changes observed at P11, only 98 differentially translated genes were identified following BH4 treatment under hypoxic conditions (62 increased and 36 decreased transcripts; Figure S1A). Correspondingly, overlap between the P30 hypoxia-responsive transcript sets was limited, with only nine hypoxia-suppressed transcripts restored by BH4 and 21 hypoxia-induced transcripts reduced following treatment (Figure S1B). Functional enrichment analysis of these residual BH4-responsive transcripts identified a restricted set of molecular functions, including oxidoreductase activity among transcripts increased by BH4 and ion transport-and membrane-associated functions among transcripts reduced by BH4 (Figure S1C, D).

Together, our differential translatome analyses demonstrate that BH4 supplementation preferentially restores ribosome-associated transcripts suppressed by developmental hypoxia during the critical period of OL maturation, while simultaneously normalizing a smaller subset of hypoxia-induced transcripts. The markedly reduced translational response observed at P30 suggests that BH4-mediated translatome remodeling is largely restricted to the earlier developmental window.

### Network analysis identifies reciprocal translational programs underlying BH4-mediated rescue

While differential translation analysis identified thousands of hypoxia- and BH4-responsive transcripts, these analyses do not determine whether translational changes occur as coordinated biological programs. To identify co-regulated ribosome-associated gene networks associated with developmental hypoxia and BH4 treatment, we performed weighted gene co-expression network analysis (WGCNA) across all experimental groups (Figure 2A). Correlation of module eigengenes with experimental condition identified two modules exhibiting reciprocal relationships with hypoxia and BH4 treatment. Module #1 was significantly reduced during developmental hypoxia and partially restored following BH4 supplementation, whereas Module #3 was increased during hypoxia and shifted toward normoxic levels after BH4 treatment (Figure 2B, C).

**Figure 2.**
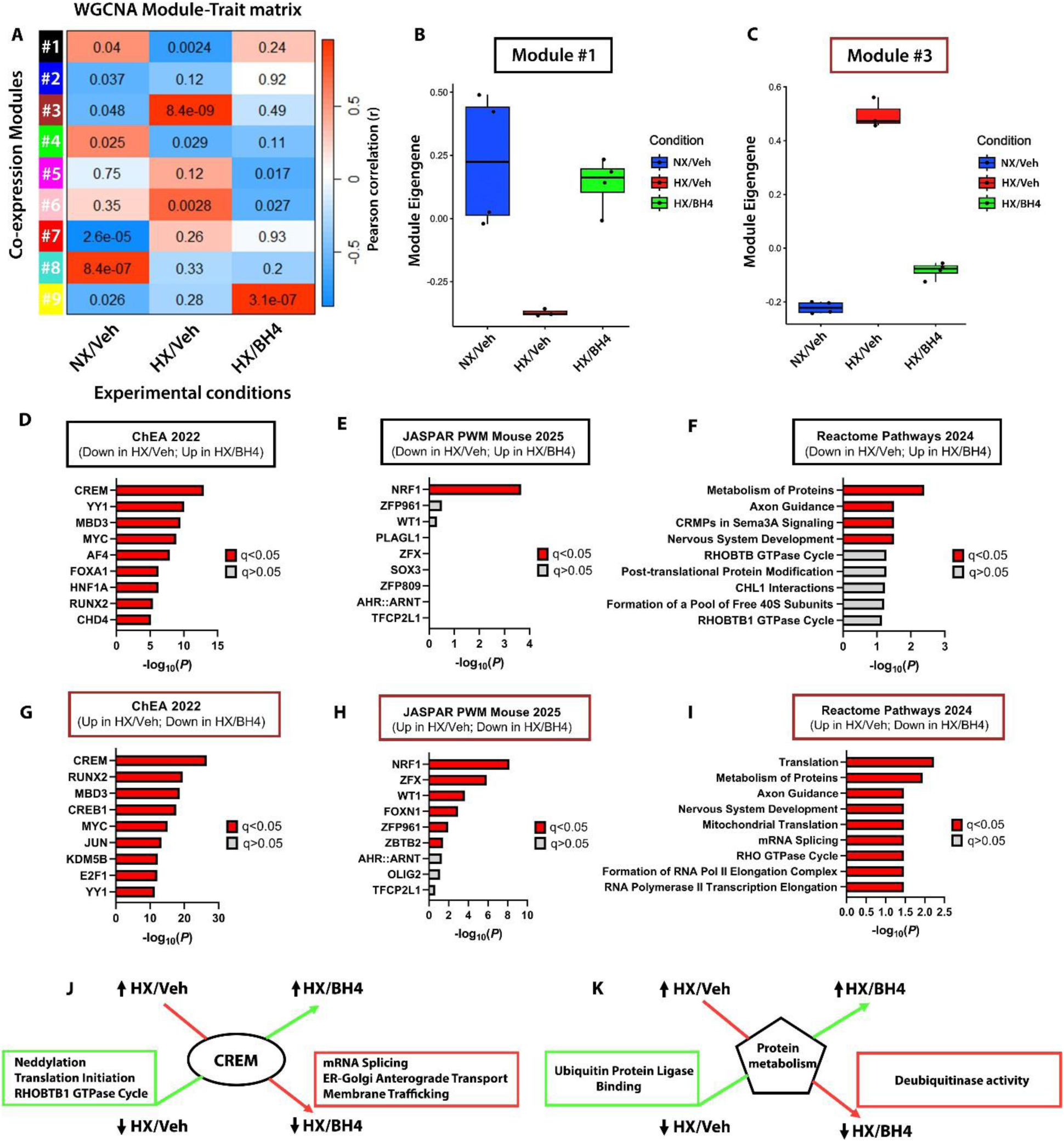
Co-expression network analysis reveals reciprocal regulation of functionally related translational programs during hypoxia and BH4-mediated rescue. (A) Weighted gene co-expression network analysis (WGCNA) module–trait correlation matrix. Each row represents an expression module, and each column corresponds to an experimental condition (NX/Veh, HX/Veh, HX/BH4). Color scale indicates Pearson correlation coefficients between module eigengenes and treatment groups, with associated P values shown within each cell. (B–C) Module eigengene expression for representative BH4-responsive modules. Module #1 (B) is negatively associated with hypoxia and restored by BH4, whereas Module #3 (C) displays reciprocal regulation, suggesting coordinated redistribution of ribosome-associated transcripts across distinct translational programs. Individual points represent biological replicates and boxplots indicate median and interquartile range. (D–I) Functional enrichment analyses of representative WGCNA modules. Panels D–F summarize transcription factor, DNA binding motif, and Reactome pathway enrichment for transcripts reduced by hypoxia and restored by BH4 (Module #1). Panels G–I show complementary enrichment analyses for transcripts increased by hypoxia and reduced following BH4 treatment (Module #3). Despite opposite patterns of regulation, both modules converge on related biological processes including protein metabolism, RNA processing, intracellular transport, and translational regulation. (J) Conceptual summary of Module #1 (down in HX/Veh, up in HX/BH4). This module is enriched for transcripts involved in translation initiation, mRNA splicing, ER-to-Golgi transport, membrane trafficking, RHOBTB1 GTPase signaling, and neddylation. These transcripts are preferentially associated with ribosomes under normoxic conditions, suppressed during hypoxia, and restored following BH4 treatment. (K) Conceptual summary of Module #3 (up in HX/Veh, down in HX/BH4). This reciprocally regulated module is enriched for protein metabolism, ubiquitin ligase binding, substrate binding, and deubiquitinase activity, illustrating selective translational redistribution of functionally related transcripts during hypoxia that is normalized by BH4. Correlation analyses were performed using WGCNA, and pathway enrichment analyses were conducted using ChEA, JASPAR, and Reactome databases in Enrichr. Bars in D-I denote –log10(p) values; filled red bars represent ontology categories with q<0.05.

To define the biological functions represented within these reciprocal modules, we performed transcription factor and pathway enrichment analyses. Module #1, comprising transcripts reduced by hypoxia and restored by BH4, was enriched for transcriptional regulators implicated in developmental and differentiation programs, including CREM, MYC, YY1, and RUNX2. YY1 and CREM have established roles in OL lineage regulation and myelin-associated transcriptional programs^27–30^, whereas MYC and RUNX2 are broader developmental regulators involved in cellular differentiation and morphogenetic processes^31,32^. Functional enrichment further identified axon guidance, nervous system development, post-translational modification, and RHO GTPase signaling pathways (Figure 2D–F), consistent with restoration of developmental and cytoskeletal programs disrupted by hypoxia.

Unexpectedly, Module #3, comprising transcripts increased during hypoxia and reduced following BH4 treatment, exhibited enrichment for many of the same broad functional domains despite exhibiting an opposing pattern of ribosome association. CREM again emerged among the top predicted upstream regulators, together with RUNX2, MBD3, JUN, and E2F1, while pathway analysis revealed enrichment for RNA processing and proteostatic programs, including RNA metabolism, mRNA splicing, translation, and RNA polymerase II–dependent transcription (Figure 2G–I), indicating a reciprocal regulation of core gene expression machinery rather than pathway exclusivity. In addition to these reciprocal modules, WGCNA identified Module #9 as being selectively associated with BH4 treatment (Figure S2A). Unlike Modules #1 and #3,

Module #9 was not principally defined by hypoxic regulation but instead represented a BH4-associated translational and developmental activation program. Upstream regulator analysis identified NRF1 together with CREM, OLIG2, ZFP961, and ZBTB2 as the strongest predicted transcriptional regulators (Figure S2B, C), while biological process enrichment highlighted neuronal developmental and metabolic programs, including axonogenesis, neuronal projection morphogenesis, neuronal differentiation, and amino acid transport (Figure S2D), consistent with activation of a BH4-responsive growth and differentiation state.

Although Modules #1 and #3 exhibited opposing patterns of ribosome association, comparison of their enriched biological processes revealed substantial functional convergence on protein homeostasis. However, this convergence reflected distinct components of the proteostasis network. Module #1 was enriched for processes associated with protein synthesis and trafficking, including translation initiation, ER-to-Golgi transport, membrane trafficking, and RHO-family GTPase signaling, whereas Module #3 preferentially captured protein turnover and RNA regulatory functions, including ubiquitin ligase binding, deubiquitinase activity, substrate recognition, and broader RNA/protein metabolic control (Figure 2J, K), suggesting reciprocal modulation of protein production versus protein degradation and RNA processing machinery.

These findings demonstrate that developmental hypoxia redistributes ribosome association across coordinated functional networks, while BH4 rebalances translational output toward OL maturation-associated programs and away from hypoxia-associated protein quality-control pathways. These transcriptomic changes are consistent with our previous in vivo observations that BH4 promotes OL maturation and improves developmental myelination following neonatal hypoxia^18^, providing a mechanistic framework for the previously observed phenotypic rescue.

### BH4 rebalances ribosome-associated gene expression toward OL maturation programs

To further define the biological programs preferentially restored by BH4, we performed targeted functional analyses on transcript subsets associated with OL maturation and translation. Candidate gene sets were generated by intersecting hypoxia-responsive, BH4-rescued differentially translated genes identified by DESeq2 with independently curated Gene Expression Omnibus (GEO)-derived gene lists linked to OL maturation or translation-associated processes (Figure 3A). Heatmap visualization demonstrated reciprocal regulation of these transcript sets across experimental groups, with developmental hypoxia reducing ribosome association of OL maturation-associated transcripts while increasing ribosome association of translation initiation-associated transcripts. BH4 treatment shifted both gene sets toward the normoxic profile, restoring translation of OL-associated transcripts while attenuating the hypoxia-induced translation signature (Figure 3A).

**Figure 3.**
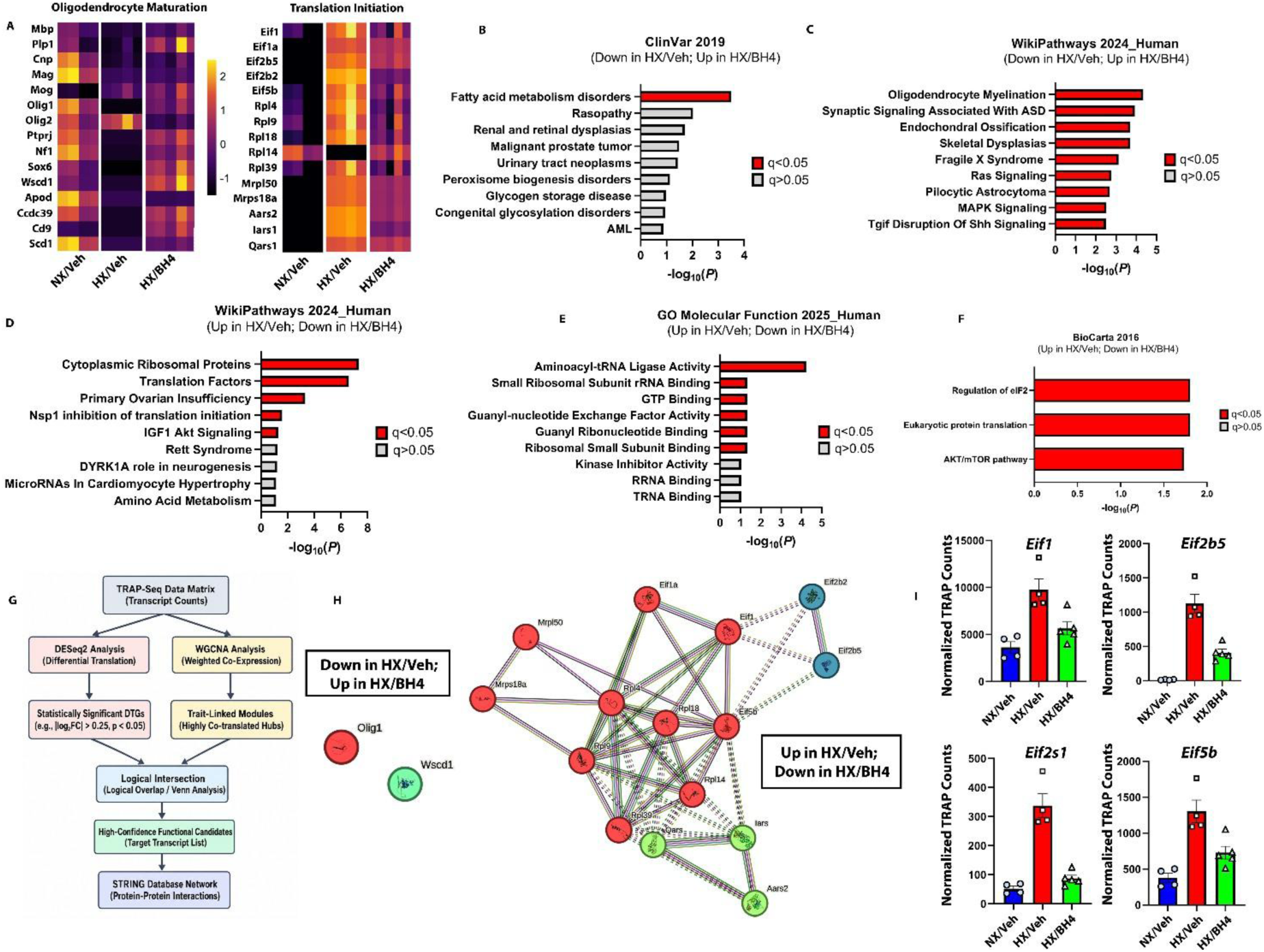
BH4 rebalances OL translational programs toward maturation following developmental hypoxia. (A) Heatmaps of representative transcripts associated with OL maturation (left) and translation initiation (right) across experimental groups (NX/Veh, HX/Veh, and HX/BH4). Z-scored ribosomal enrichment values illustrate hypoxia-induced suppression of multiple maturation-associated transcripts and their partial restoration following BH4 treatment. (B–F) Functional enrichment analyses of BH4-responsive transcripts. Gene ontology and pathway analyses demonstrate that BH4 preferentially restores ribosome-associated transcripts involved in OL maturation, myelination, lipid metabolism, and protein translation, while reducing enrichment of hypoxia-associated stress and protein homeostasis programs. Together these analyses indicate a shift in translational output toward developmental maturation. (G) Bioinformatic workflow for identifying high-confidence BH4-responsive transcripts. Differentially translated genes (DESeq2) were intersected with WGCNA module genes exhibiting reciprocal regulation by hypoxia and BH4 to generate high-confidence transcript sets for protein interaction network analysis. (H) STRING protein interaction networks of high-confidence BH4-responsive transcripts. Left, transcripts reduced by hypoxia and restored by BH4 identified only two interconnected OL maturation genes (*Olig1* and *Wscd1*), precluding meaningful network analysis. Right, transcripts increased by hypoxia and reduced by BH4 formed a highly connected network enriched for translation initiation factors and associated ribosomal proteins. Node colors denote functional clusters identified by k-means clustering (*k* = 3), highlighting distinct but interconnected translation-related subnetworks. (I) Normalized TRAP-seq read counts for representative translation initiation genes (*Eif1, Eif2b5, Eif2s1, and Eif5b*) identified from the overlapping DESeq2/WGCNA analysis. These transcripts were elevated during hypoxia and reduced following BH4 treatment, validating the expression patterns observed in the heatmap and network analyses. Bars represent mean ± SEM with individual biological replicates. Pathway enrichment analyses were performed using WikiPathways, Reactome, Gene Ontology, BioCarta, ClinVar, and ChEA databases in Enrichr. Bars in B-F denote – log10(p) values; filled red bars represent ontology categories with q<0.05. Protein interaction networks were generated using STRING, with functional clusters assigned by k-means clustering (*k* = 3). Statistical significance was defined as adjusted P < 0.05.

To characterize the biological functions represented within the OL maturation-associated transcript set, enrichment analyses were performed using Enrichr. ClinVar disease enrichment identified fatty acid metabolism disorders as the highest-ranking shared disease category among transcripts suppressed by hypoxia and restored following BH4 treatment (Figure 3B). Consistent with this finding, WikiPathways analysis demonstrated significant enrichment for OL maturation leading to myelin formation, together with Ras signaling, synaptic signaling, and additional pathways involved in nervous system development and glial maturation (Figure 3C). These findings support selective restoration of translational programs required for OL maturation rather than generalized increases in ribosome-associated translation.

We next examined transcripts exhibiting the reciprocal pattern of regulation, comprising genes increased during hypoxia and reduced following BH4 treatment. Enrichment analyses demonstrated significant overrepresentation of biological processes associated with translation, translational initiation, RNA metabolism, peptide biosynthesis, ribosome biogenesis, and protein metabolism (Figure 3D–F), indicating that developmental hypoxia preferentially increases ribosome association of genes involved in translational machinery and protein homeostasis, which are subsequently normalized by BH4 treatment.

We then focused on the subset of OL maturation-associated transcripts that reached differential enrichment significance across both DESeq2 comparisons. Volcano plots confirmed that developmental hypoxia suppressed ribosome association of these maturation-associated genes, whereas BH4 treatment restored their ribosome association (Figure S3A, B). To investigate upstream regulatory mechanisms associated with this BH4-responsive transcript subset, ChEA enrichment analysis identified CREM as the highest-ranked predicted transcriptional regulator (Figure S3C). Gene Ontology molecular function analysis further demonstrated enrichment for protein tyrosine phosphatase activity, PDGFR binding, and histone acetyltransferase binding (Figure S3D), indicating enrichment of receptor-mediated signaling and chromatin regulatory pathways within the BH4-responsive transcript subset. Since multiple analyses converged on translation initiation and eIF2-associated pathways, we next examined phosphorylation of eIF2α, a central regulator of translation initiation whose phosphorylation suppresses canonical cap-dependent protein synthesis while promoting selective translation of stress-responsive transcripts. Immunofluorescence demonstrated a significant reduction in the p-eIF2α/eIF2α ratio within OLIG2-positive OLs following developmental hypoxia (Figure S3E, F), consistent with altered regulation of eIF2α-dependent translation initiation at P11. BH4 supplementation did not significantly restore eIF2α phosphorylation at the P11 time point examined. These findings provide independent in vivo evidence that developmental hypoxia alters a major regulatory pathway controlling translation initiation, consistent with the enrichment of translation initiation and eIF2-associated transcripts identified by TRAP-seq. Together with the TRAP-seq analyses, these data suggest that BH4-mediated translational rescue occurs without normalization of steady-state eIF2α phosphorylation at this developmental stage. To prioritize the most robust candidate regulators within these reciprocal programs, differentially translated genes identified by DESeq2 were intersected with BH4-responsive WGCNA modules to generate high-confidence transcript sets for STRING protein interaction analysis (Figure 3G). Among transcripts reduced by hypoxia and restored following BH4 treatment, only *Olig1* and *Wscd1* were identified by both analytical approaches, precluding construction of a meaningful interaction network but highlighting these genes as high-confidence candidates associated with BH4-mediated restoration of OL maturation (Figure 3H, left). In contrast, transcripts increased during hypoxia and reduced by BH4 formed a highly interconnected protein interaction network dominated by translation initiation factors and translational regulators. K-means clustering resolved this network into three functional clusters corresponding to large ribosomal subunit components (red), aminoacyl-tRNA biosynthesis (green), and recycling of eIF2:GDP (blue) (Figure 3H, right), indicating coordinated regulation of multiple stages of translational initiation and protein synthesis during developmental hypoxia. To validate these network findings, normalized TRAP-seq read counts were examined for representative translation initiation factors identified through the integrated DESeq2/WGCNA analysis. *Eif1*, *Eif2b5*, *Eif2s1*, and *Eif5b* each exhibited increased ribosome association during developmental hypoxia that was consistently attenuated by BH4 treatment (Figure 3I).

### BH4 restores translational regulation and nuclear localization of the OL lineage regulator OLIG1

Integrated differential translation and network analyses identified *Olig1* as one of only two OL maturation-associated transcripts consistently prioritized by both DESeq2 and WGCNA analyses (Figure 3H), prompting further investigation of its regulation across multiple molecular layers. Because TRAP-seq measures ribosome-associated mRNA rather than total transcript abundance, we next asked whether BH4-mediated rescue of *Olig1* reflects effects on translational engagement, transcript abundance, or both.

Normalized TRAP-seq read counts demonstrated that developmental hypoxia significantly reduced ribosome association of *Olig1*, whereas BH4 treatment restored ribosome occupancy toward normoxic levels (Figure 4A), indicating a strong effect of BH4 on translational engagement of this transcript. To determine whether these changes were accompanied by alterations in total transcript abundance, RNAscope in situ hybridization was performed within the white matter. Consistent with a hypoxia-induced reduction in *Olig1* expression at the mRNA level, quantification of *Olig1* transcript puncta signal intensity within OL lineage cells demonstrated a significant decrease following developmental hypoxia. However, in contrast to the robust recovery observed in TRAP-seq, BH4 treatment did not significantly restore *Olig1* mRNA abundance (Figure 4B, C), indicating that BH4 effects on *Olig1* are not mediated through normalization of transcript levels and instead selectively impact ribosome association.

**Figure 4.**
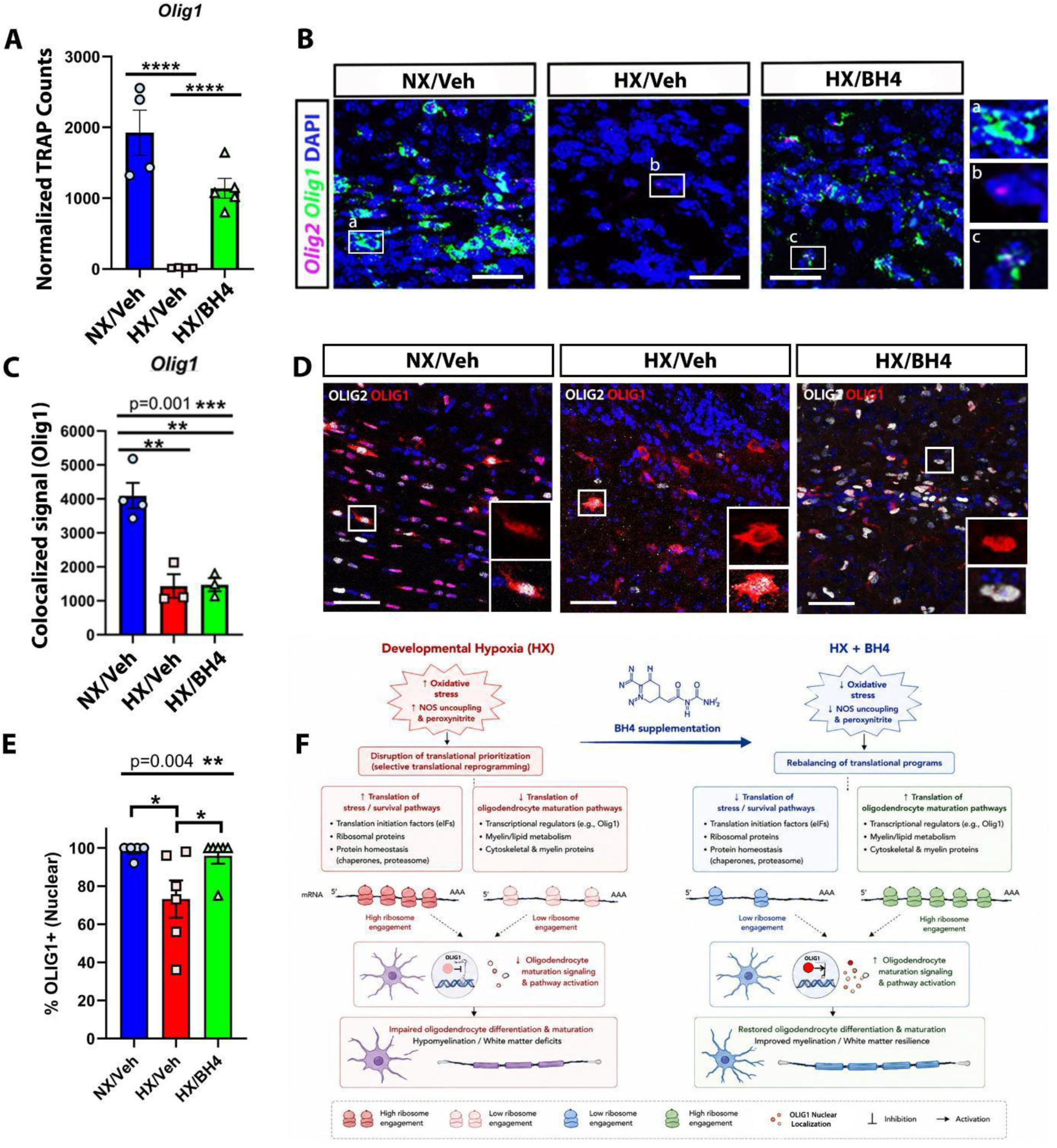
Translational restoration of *Olig1* by BH4 is accompanied by recovery of OLIG1 protein nuclear localization. (A) Normalized TRAP-seq read counts for *Olig1*. Developmental hypoxia markedly reduced ribosome-associated *Olig1* transcripts, whereas BH4 significantly restored ribosomal enrichment toward normoxic levels. (B) Representative RNAscope images depicting *Olig1* transcript expression in white matter tracts from NX/Veh, HX/Veh, and HX/BH4 mice. (C) Quantification of *Olig1* transcript puncta expression within OL lineage cells, measured as *Olig1/Olig2* colocalized fluorescence intensity. Hypoxia reduced *Olig1* transcript expression, and BH4 treatment did not restore overall transcript levels. (D) Representative immunofluorescence images showing OLIG1 (red), OLIG2 (white), and merged images across treatment groups. Insets highlight representative OLs illustrating reduced OLIG1 nuclear localization following hypoxia and its recovery following BH4 treatment. (E) Quantification of OLIG1-positive nuclei among OL lineage cells. Developmental hypoxia significantly reduced nuclear localization of OLIG1, while BH4 restored the proportion of OLIG1-positive nuclei toward normoxic levels. (F) Proposed mechanism of BH4-mediated restoration of OL maturation during developmental hypoxia. Data are presented as mean ± SEM with individual biological replicates. Statistical significance was determined using one-way ANOVA followed by Tukey’s multiple-comparison testing for panels A and C, and the Kruskal– Wallis test followed by Dunn’s multiple-comparison test for panel E (* P < 0.05, ** P < 0.01, *** P < 0.001, **** P < 0.0001). Scale bar in B = 50 µm; D = 100 µm.

We next examined whether these differences in transcript abundance and ribosome engagement were accompanied by alterations in OLIG1 protein localization. Nuclear localization of OLIG1 is required for OL maturation and activation of myelin-associated gene expression. Immunofluorescence revealed a marked reduction in nuclear OLIG1 signal within OLIG2-positive OL lineage cells following developmental hypoxia. In contrast, BH4 treatment restored nuclear OLIG1 localization toward normoxic levels (Figure 4D, E).

Together, these findings demonstrate that developmental hypoxia disrupts *Olig1* regulation at multiple molecular levels, including transcript abundance, ribosome association, and nuclear accumulation of OLIG1 protein. BH4 selectively restores ribosome association and nuclear localization despite persistent reductions in transcript abundance, identifying OLIG1 as a candidate mediator of BH4-dependent translational rescue during OL maturation (Figure 4F), thereby supporting translational and post-transcriptional mechanisms of OL maturation that are not fully captured by steady-state transcript abundance alone.

## DISCUSSION

Developmental white matter injury in CHD is strongly linked to chronic cerebral hypoxemia and is a major contributor to long-term neurodevelopmental impairment, yet the molecular mechanisms connecting sustained reductions in oxygen delivery to OL maturation arrest and diffuse hypomyelination remain incompletely defined. In this study, we identify translational reprogramming within postmitotic OLs as a central mechanism underlying hypoxia-induced white matter dysmaturation and demonstrate that supplementation with the NOS cofactor BH4 selectively restores ribosome association of key OL maturation transcripts. Given our data indicating that a large fraction (∼90%) of transcripts suppressed by hypoxia are restored by BH4, this study demonstrates the central role of BH4 and coupled NOS in oxidative stress-induced OL dysmaturation. These findings provide key mechanistic insight into how chronic hypoxemia disrupts white matter development in CHD while revealing translational rescue as a previously underrecognized mode of action for BH4 in the developing brain.

Consistent with prior work showing that hypoxia arrests OL maturation and leads to diffuse hypomyelination in both human infants with CHD and experimental models of chronic hypoxia^8,11,33,34^, we found that hypoxia extensively remodeled ribosome-associated gene expression within postmitotic OLs. TRAP-seq identified nearly 3,000 differentially translated genes at P11, revealing a coordinated redistribution of ribosome engagement rather than global suppression of protein synthesis. Transcripts associated with OL maturation, including *Olig1* and genes involved in Ras/MAPK signaling, cytoskeletal organization, and myelin biosynthesis, exhibited reduced ribosome association, whereas transcripts involved in translation initiation, RNA metabolism, and protein homeostasis were increased. WGCNA further demonstrated that these changes were organized into reciprocal gene networks, indicating that developmental hypoxia shifts translational output away from maturation-associated biosynthesis toward stress-responsive proteostatic programs.

A key finding of this study is that BH4 does not function as a global activator of translation but instead selectively restores ribosome association of hypoxia-suppressed transcripts. Most BH4-responsive genes overlapped with hypoxia-downregulated transcripts, and rescue efficiency was significantly higher for these genes compared with hypoxia-induced transcripts. This asymmetry indicates that BH4 preferentially restores developmental translational programs rather than broadly reversing hypoxia-induced changes in gene expression. Importantly, this effect was temporally restricted, with minimal translational rescue observed at P30, suggesting that maturing OLs exhibit heightened translational plasticity during a defined developmental window corresponding to active myelination. Mechanistically, these findings are consistent with the established role of BH4 in maintaining NOS coupling and limiting oxidative stress^14,35,36^, thereby preserving redox-sensitive pathways required for protein synthesis. Rather than globally increasing translational capacity, BH4 appears to selectively restore ribosome engagement of OL maturation-associated transcripts suppressed by hypoxia.

*Olig1* emerged as a high-confidence regulatory node across both differential translation and network analyses. Although developmental hypoxia reduced both *Olig1* transcript abundance and ribosome association, BH4 selectively restored ribosome engagement and nuclear localization of OLIG1 protein without normalizing mRNA levels, suggesting that functional recovery of OL maturation can occur through post-transcriptional mechanisms. Whether enhanced ribosome association directly contributes to OLIG1 nucleocytoplasmic shuttling remains to be determined. During OL development, nuclear localization of OLIG1 is a critical regulator of OL maturation and activation of myelin-associated gene expression. Appropriate nuclear retention of OLIG1 is required for progression through the myelinating program^37–39^. The failure of OLIG1 to enter or persist in the nucleus under hypoxic conditions aligns with known redox sensitivities of nuclear import machinery, where oxidative stress can disrupt Ran GTPase gradients or directly modify karyopherin-dependent transport^40–42^. Furthermore, our network analysis identified the cAMP-responsive element modulator (CREM) as a top predicted upstream regulator across multiple BH4-responsive networks. cAMP/PKA signaling is well-established as a potent driver of OL maturation and myelin gene expression^29,43,44^. It is highly plausible that BH4-mediated reduction of oxidative stress or restoration of NOS coupling^45^ reactivates downstream cAMP/CREM signaling cascades, which may contribute to post-translational regulation of OLIG1, potentially via phosphorylation^46^ or acetylation^37^, to alter its nuclear retention and stabilize its translation on the ribosome via specialized RNA-binding proteins, such as Quaking (QKI)^47^. Similarly, enrichment of NRF1 target genes suggests restoration of mitochondrial and metabolic biosynthetic programs required for OL maturation^48–50^.

Recent studies have established intimate links between redox signaling, NOS activity, and translational control. Oxidative stress alters ribosomal proteins, elongation factors, and mRNA-binding proteins, whereas physiological nitric oxide regulates protein synthesis through eIF2α, ribosome biogenesis, mTOR, and AMPK signaling^51–54^. Accordingly, BH4 deficiency may impair OL maturation by disrupting multiple redox-sensitive mechanisms governing translation. Consistent with altered translational regulation, hypoxic postmitotic OL cells exhibited reduced p-eIF2α/eIF2α ratios together with increased ribosome association of core translation initiation factors (*Eif1, Eif2b5, Eif2s1,* and *Eif5b*). Although acute hypoxia typically activates the integrated stress response (ISR) by increasing eIF2α phosphorylation^55,56^, chronic hypoxia may instead engage adaptive feedback pathways, including GADD34-mediated eIF2α dephosphorylation^57–59^. The accumulation of translation initiation factors may therefore represent a compensatory attempt to maintain protein synthesis under sustained stress. Notably, BH4 reduced this increased association without restoring eIF2α phosphorylation, suggesting that BH4-mediated translational rescue operates largely independently of canonical ISR signaling and instead reflects broader reorganization of ribosome engagement during chronic hypoxia^60^.

Disrupted white matter development before birth is an established contributor to neurodevelopmental impairments in children with complex CHD^61^. Our findings support a model in which redox imbalance and translational suppression converge to arrest OL maturation, while BH4 restores translation of maturation-associated transcripts required for myelination. These observations are consistent with our previous demonstration that pharmacological BH4 restoration during hypoxia improves OL maturation and myelin formation^18^ and further suggest that restoration of metabolic pathways, including Ras/MAPK signaling, and fatty acid metabolism, may alleviate the biosynthetic constraints imposed by chronic hypoxia^62,63^. Because hypoxia suppresses oxygen-dependent steps in lipid and cholesterol biosynthesis^64,65^, our findings suggest that BH4 may alleviate this biosynthetic constraint by restoring ribosome association of transcripts encoding key components of these pathways. Given the excellent safety record of BH4 in pregnancy, these findings support pharmacological BH4 restoration as a potential adjunct therapy for protecting the developing white matter in fetuses with CHD. Beyond CHD, these results may have broader relevance for perinatal brain injury contexts involving hypoxia, oxidative stress, or metabolic insufficiency, including preterm birth, neonatal encephalopathy, and placental insufficiency, where OL maturation vulnerability is a common pathological feature.

This study has several limitations. Firstly, TRAP sequencing isolates ribosome-associated mRNAs but does not directly measure protein output. Therefore, some translational changes may not correlate linearly with protein abundance. Future studies using proteomics or metabolic labeling approaches could complement these findings. Secondly, although BH4 improved transcript translation in OL lineage cells, it may also exert effects on neurons, vascular cells, or microglia, given their known dependence on redox and NOS signaling pathways. Disentangling cell-type–specific effects will require single-cell targeted omic approaches. Finally, while the period of white matter maturation from P3 to P11 in mice corresponds approximately to the third trimester of human brain development^13^, the chronic hypoxia paradigm used here primarily models reduced oxygen availability and does not capture the complex hemodynamic and cardiovascular abnormalities characteristic of fetal and neonatal CHD.

Several potential future studies emerge from this work. First, the specific molecular intermediates linking BH4 to translational rescue need to be delineated. Candidate mechanisms include modulation of NOS-dependent cGMP signaling, mitigation of oxidative ribosomal damage, and normalization of mTORC1 pathways. Second, future studies should examine whether BH4 enhances translation by altering tRNA charging, ribosomal biogenesis, or translation factor activity. Third, combining BH4 supplementation with metabolic interventions that support mitochondrial function may yield synergistic benefits for OL maturation in hypoxic environments. Finally, while BH4 supplementation has been shown to protect against fetal hypoxic-ischemic injury in rabbit models^66,67^, establishing efficacy in large animals is a critical step toward clinical translation, particularly given that clinical trials based on rodent studies often fail especially those related to brain function^68^.

This study demonstrates that developmental hypoxia disrupts OL maturation through coordinated remodeling of the OL translatome and that pharmacological restoration of BH4 selectively restores translation of key maturation-associated transcripts. These findings identify translational rescue as a central mechanism by which BH4 mitigates white matter dysmaturation and suggest that modulation of redox-sensitive translational pathways represents a promising therapeutic strategy for protecting the developing brain in fetuses and infants with CHD.

