## Supplementary Information for "Tetrahydrobiopterin restores oligodendrocyte maturation after hypoxic white matter injury by reprogramming translational output"

**Figure S1**

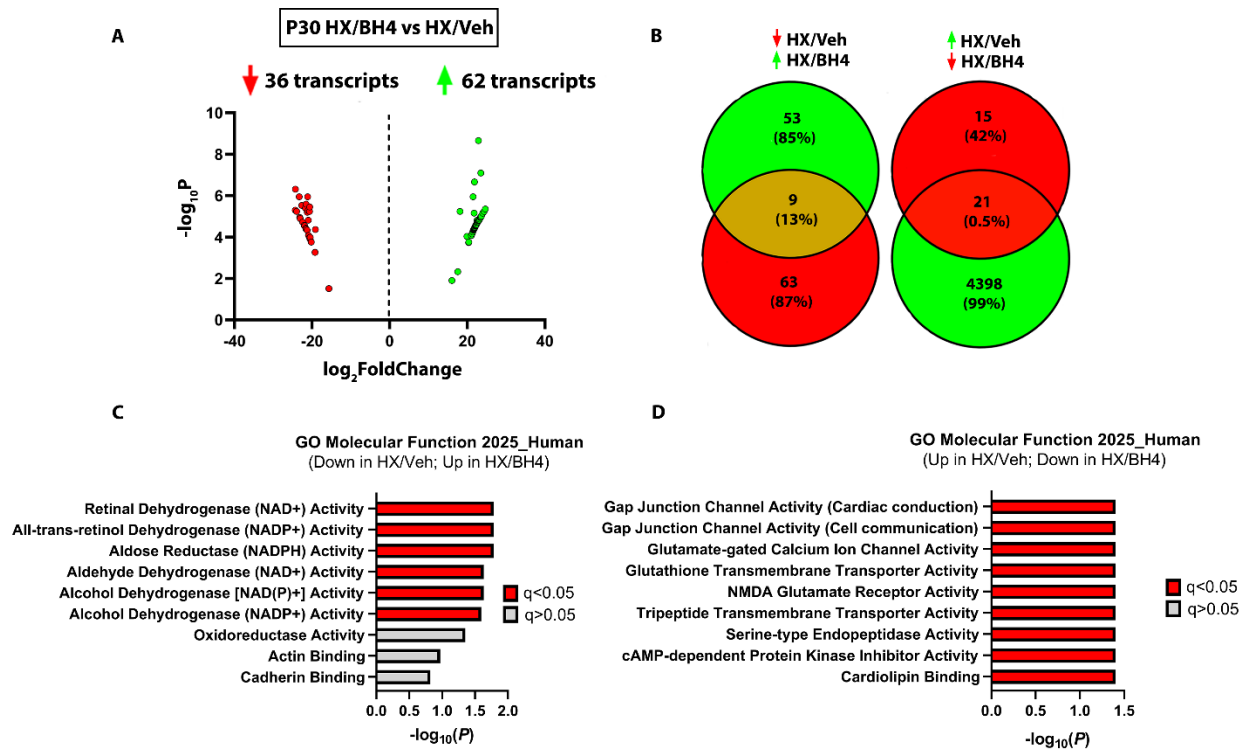

**Figure S1. Translatome analysis and Gene Ontology (GO) molecular functions of differentially translated transcripts in P30 HX/BH4 versus HX/Veh groups.** (A) Volcano plot illustrating differentially translated transcripts between P30 HX/BH4 and HX/Veh conditions. A total of 36 transcripts were significantly downregulated (red dots/arrow) and 62 transcripts were significantly upregulated (green dots/arrow) in their ribosomal enrichment. (B) Venn diagrams illustrating overlap between hypoxia-responsive DTGs and BH4-responsive DTGs at P30. Left, transcripts downregulated by hypoxia and restored by BH4. Right, transcripts upregulated by hypoxia and reduced by BH4. Central intersections represent BH4-rescued translational targets. (C-D) Molecular function enrichment analyses of P30 BH4-responsive DTGs. Panel C shows enrichment analysis for transcripts decreased by hypoxia and increased following BH4 treatment. Panel D shows complementary enrichment analysis for transcripts increased by hypoxia

and reduced following BH4 treatment. Enrichr. Bars in C-D denote $-\log_{10}(p)$ values; filled red bars represent ontology categories with $q < 0.05$.

**Figure S2**

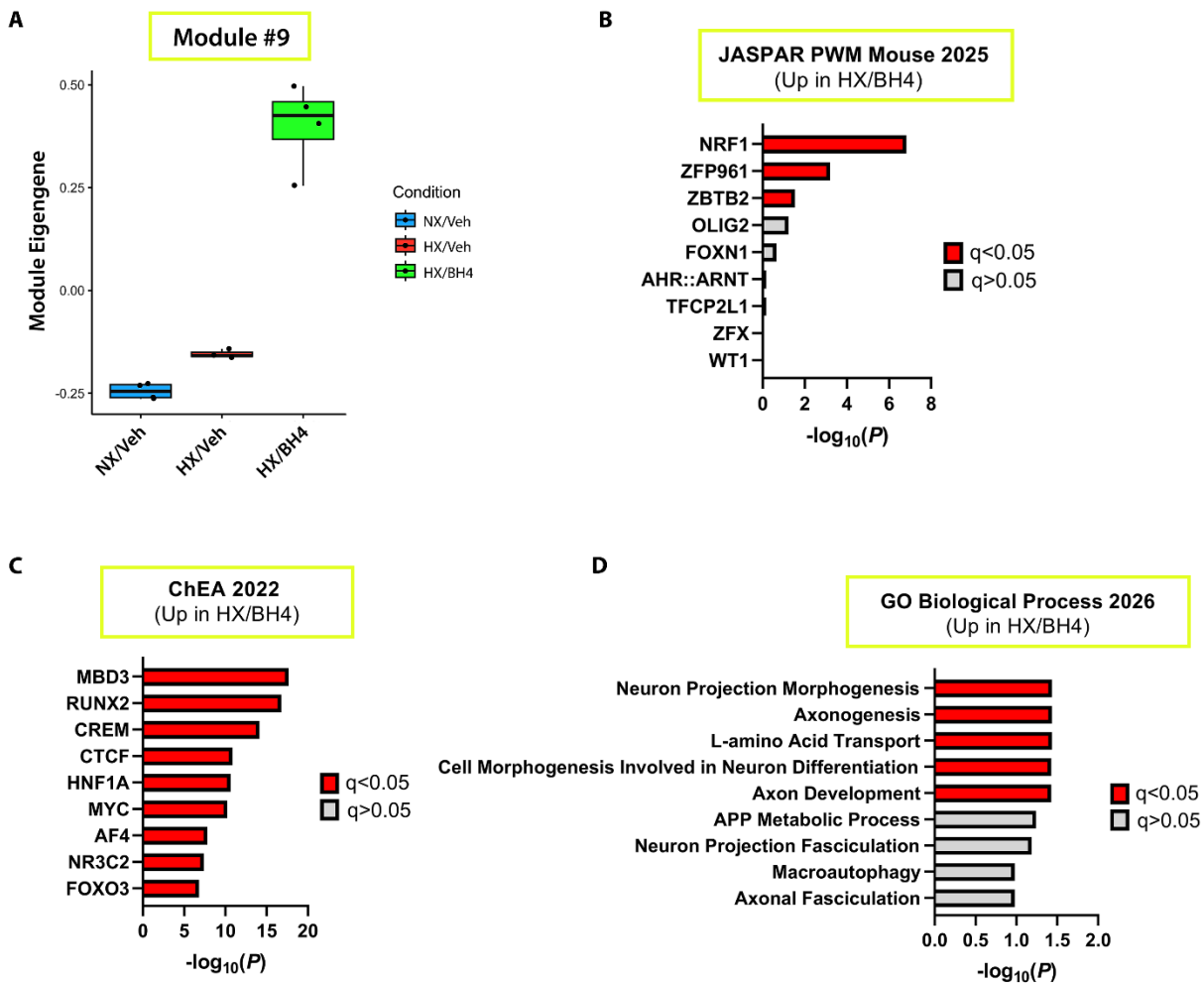

**Figure S2. WGCNA identifies an upregulated translational module unique to BH4 treated hypoxia and enriched for neuronal development and CREM/NRF1 regulatory programs.** (A) Module eigengene expression for BH4-upregulated module #9. Individual points represent biological replicates and boxplots indicate median and interquartile range. (B-C) JASPAR PWM and ChEA 2022 transcription factor motif enrichment analysis of Module #9 transcripts. Predicted upstream regulators include

NRF1, ZFP961 and ZBTB2, MBD3 and CREM, suggesting coordinated regulation of BH4-responsive translational programs involved in neural development and cellular metabolism. (D) Gene Ontology Biological Process enrichment analysis of Module #9 demonstrating significant enrichment for axonogenesis, neuron projection morphogenesis, neuronal differentiation, regulation of nervous system development, amino acid transport, and related developmental pathways. Bars in B-D denote $-\log_{10}(p)$ values; filled red bars represent ontology categories with $q < 0.05$.

**Figure S3**

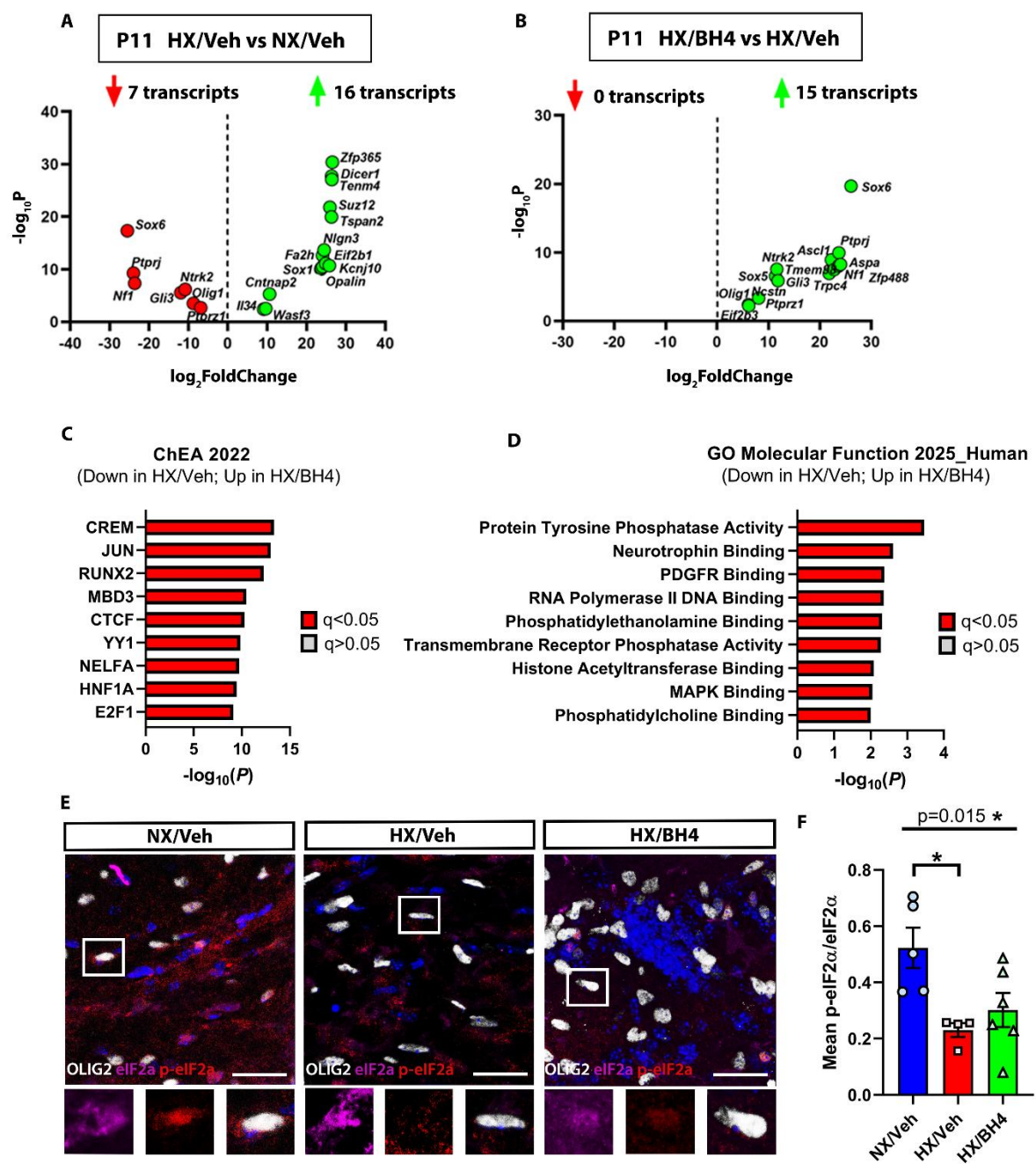

**Figure S3. Additional validation of BH4-responsive translational programs and EIF2 signaling following developmental hypoxia.** (A) Volcano plot of differentially translated genes (DTGs) in HX/Veh versus NX/Veh at P11 highlighting representative oligodendrocyte maturation-associated transcripts identified by TRAP-seq. Red points

indicate transcripts significantly reduced by hypoxia and green points indicate transcripts increased by hypoxia ($|\log_2FC| \geq 0.25$, adjusted $P < 0.05$). (B) Volcano plot of DTGs in HX/BH4 versus HX/Veh demonstrating restoration of selected hypoxia-responsive transcripts following BH4 treatment. Representative genes are annotated. Red points indicate transcripts reduced by BH4 and green points indicate transcripts increased by BH4 ($|\log_2FC| \geq 0.25$, adjusted $P < 0.05$). (C) ChEA 2022 transcription factor enrichment analysis of transcripts downregulated by hypoxia and restored by BH4. (D) Gene Ontology (Molecular Function 2025) enrichment analysis of transcripts increased by hypoxia and reduced by BH4. (E) Representative immunofluorescence images and quantification (F) of p-eIF2α (red) expression (normalized to total eIF2α (magenta)) within OLIG2-positive oligodendrocyte lineage cells (white) in the white matter of NX/Veh, HX/Veh, and HX/BH4 mice. DAPI (blue) labels nuclei. Insets show higher-magnification views of representative cells. Note that chronic hypoxia results in significantly reduced p-eIF2α, which is not restored by BH4 supplementation. Bar graphs represent mean ± SEM with individual biological replicates. Bars in C-D denote $-\log_{10}(p)$ values; filled red bars represent ontology categories with $q < 0.05$. Statistical significance for E-F was determined using one-way ANOVA with Tukey's multiple-comparison testing (\* $P < 0.05$). Scale bar = 50 $\mu\text{m}$.
